# Reporting quality of trend analyses published in leading medicine and oncology journals during 2008-2018

**DOI:** 10.1101/2020.09.18.303701

**Authors:** Xiaoling Yuan, Fei Deng, Yating Wang, Lanjing Zhang

## Abstract

Reporting quality of clinical research is critical for evidence-based medicine and reproducibility of clinical research. Most of the works focused on reporting quality of clinical trials and observational longitudinal studies. However, few focused on that of trend analyses. The reporting of recommended statistic metrics in trend analyses was also largely unclear. Therefore, we examined reporting quality of the trend analyses based on reporting of recommended statistic metrics. We systemically searched the PubMed for the trend-analysis articles published in 10 leading medicine and oncology journals during the 11 years from 2008 to 2018. The studies published after 2019 were not included due to the sudden, significant increase of publication number during and immediately after the COVID-19 pandemic. Only original articles, research letters and meta-analyses/systematic reviews were included. We scored the reporting quality of these articles based on whether they reported p-values/effect-sizes, and beta/co-efficient/slope/annual-percentage-change (APC). There were 297 qualified articles, among which 193 (66.0%) and 216 (72.7%) articles reported P-value and effect-size, respectively. Only 13 (5.8%) analyses reported neither p-value/effect size nor beta/coefficient/slope/APC. In multivariable regression models, author affiliation of epidemiology department was associated with less reporting effect-size, but that of statistics department with more reporting. Interestingly, U.S. senior-authors (versus non-U.S.) more likely reported p-values. No factors were independently linked to reporting APC. The reporting quality of trend analyses in leading medicine and oncology journals appear moderate and should be further improved. We thus call for more research and awareness of reporting-quality in trend analyses in oncology research and beyond.

## Introduction

Reporting quality of clinical research is critical for evidence-based medicine and reproducibility of clinical research. Most of the works focused on reporting quality of clinical trials, including those in neuro-oncology, urology, cardiology, nephrology, pharmaceutics and infectious disease.^1-7^Although the research quality of clinical trials was improving,^8,9^ that of observational longitudinal studies remains low.^10-12^ However, few studies concerned the reporting quality of trend analyses.

Trend analyses are critical for assessing changes and predicting future of epidemiological parameters.^13 14^ Recent trend-analysis guidelines recommended reporting slopes or beta/coefficient if possible.^15^ American Statistical Association and others recommended reporting effect-size.^16,17^ However, reporting of these recommended statistic metrics in trend analyses was largely unclear. Therefore, we examined reporting quality of the trend analyses in leading medicine and oncology journals, with reported p-value, effect-size or beta/coefficient/Annual percent change (APC) as the quality metrics. We also identified factors associated with reporting quality in these trend analyses.

## Methods

We systemically searched the PubMed for the articles published during the 11 years from January 1, 2008 to December 31, 2018, whose titles included “trend” or “trends” among the following medicine and oncology journals: *Ann Intern Med, Ann Oncol, BMJ, J Clin Oncol, J Natl Cancer Inst, JAMA Oncol, JAMA, Lancet, Lancet Oncol and N Engl J Med*. It was considered to include articles published after 2019. However, there were surges in publication number during the pandemic of COVID-19 (also known as SARS-CoV-2 coronavirus) and after.^18 19^ We felt the reporting quality might be influenced by the pandemic and surges in publication number. Therefore, we remain focused on the 11 years of 2008-2018.

We only included original articles, research letters and meta-analyses/systematic reviews of trend analysis. We limited our search to title words to ensure articles focused primarily on trend analysis. We acknowledge this approach might miss some relevant articles and discuss this limitation. Three authors independently reviewed full-texts using a standardized data extraction form: publication year, journal specialty (medicine/oncology), model type, reporting of *P-value*, effect-size (defined as quartiles/confidence/credible/uncertainty intervals) and co-efficient/slope/Annual Percent Change (APC), senior author(s)’ location, and any author in School of Public Health, statistics department, or epidemiology department. When there was discrepancy, they would discuss and reconcile the differences. In the rare events that a consensus could not be reached, Dr Zhang would make the final decision.

According to guidelines,^15^ co-efficient/slope/APC should be reported in (piece-wise) linear models. We assessed reporting whether co-efficient/slope/APC was reported in the articles using linear model. We also scored the reporting quality of these articles by assigning 1 point for reporting a p-value or effect-size, and another point for reporting a beta/co-efficient/slope/APC. For articles reporting the same analysis, each article would be assessed independently. The sum of each article’s scores was considered as its reporting-quality score, which was up to 2 points. Points/scores were unweighted since the metrics cover different aspects of the statistics. The clinical significance of various metrics was to our knowledge not differentiated or compared by known recommendations,^15-17^ although they are all clinically important and useful.

We used Chi-square test, Fisher exact test, and (ordinal) logistic regression to examine potential associations (Stata, version 15). Only the factors with P<0.10 shown in univariate analysis were included in multivariable logistic regression analyses. Two-sided P-values were reported. Statistical significance was considered when P<0.05.

## Results and Discussion

Among the 398 identified reports of trend analysis published during 2008-2018, there were 297 qualified reports (**Supplementary Figure**), including 38 (12.8%) analyses using non-parametric model, 226 (76.1%) analyses using (piece-wise) linear model, 32 (10.8%) analyses using non-linear parametric model and 1 (0.3%) analysis using semi-parametric model (Cox regression). Among these analyses, 193 (66.0%) and 216 (72.7%) analyses reported P-value and effect-size, respectively. Subgroup analyses showed that U.S. senior authors more likely reported P-value or effect-size than non-U.S. senior authors (**Figure 1**), while reporting of these parameters was not linked to any of the other factors.

**Figure 1.**
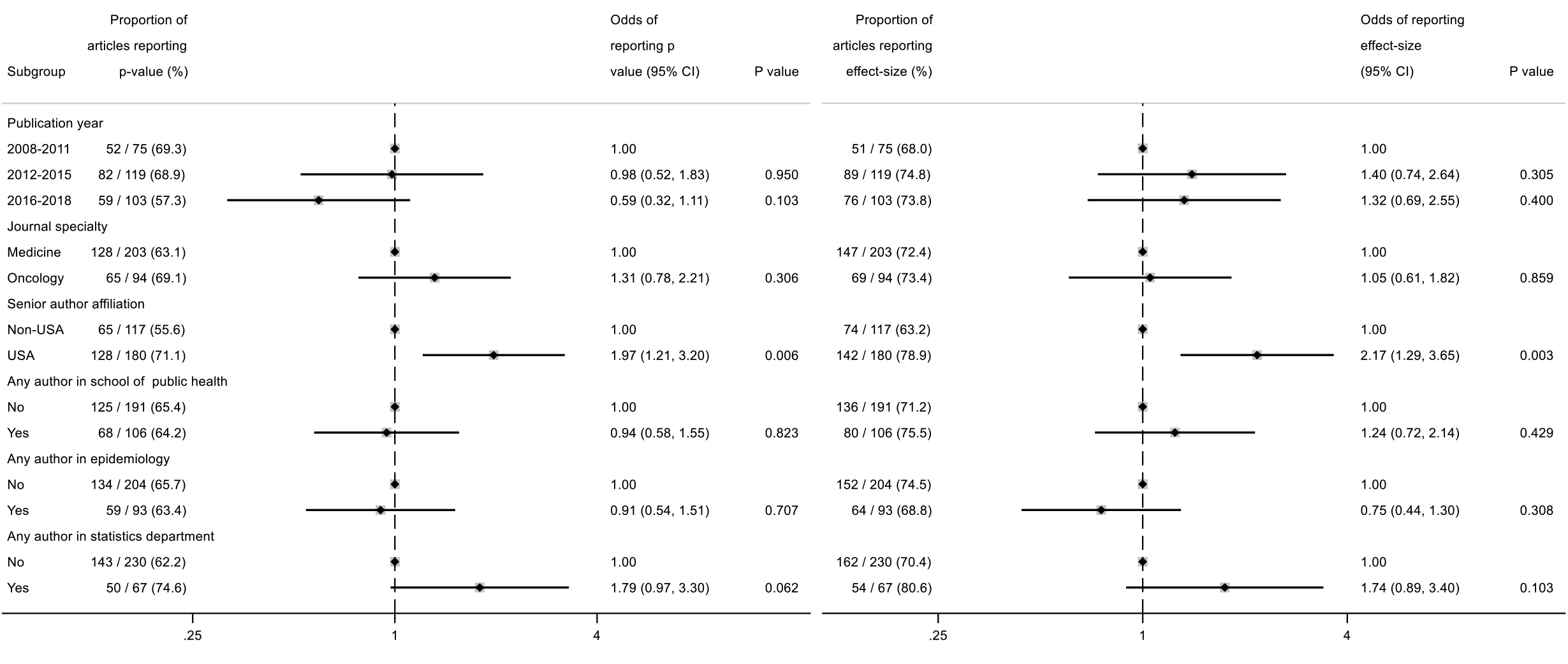
Proportions of the trend analyses published during 2008-2018 reporting p-value and effect-size and their associated factors. Only senior author’s affiliation (U.S.A. vs non-U.S.A.) was linked to reporting either p-value (Left) or effect size (Right) among the 397 trend analyses published in leading medicine and oncology journals during 2008-2018.

Among the 226 trend analyses using (piecewise) linear model (**Table 1**), 169 (74.8%) reported p-value, 183 (81.0%) reported effect-size, 94 (41.6%) reported APC and 34 (15.0%) reported Beta/coefficient/slope. No multiple articles reported the same or similar analysis. Only 13 (5.8%) analyses reported neither p-value/effect size nor beta/coefficient/slope/APC. Ordinal logistic regression showed only author affiliation with school of public health was linked to higher reporting-quality scores (Odds ratio=7.44, 95% confidence interval, 3.22 - 31.17). In multivariable regression models (**Table 2**), author affiliation with epidemiology or statistics department was associated with reporting effect-size, and U.S. senior-authors (versus non-U.S.) more likely reported p-value. No factors were independently linked to reporting APC (**Table 2**).

**Table 1.**
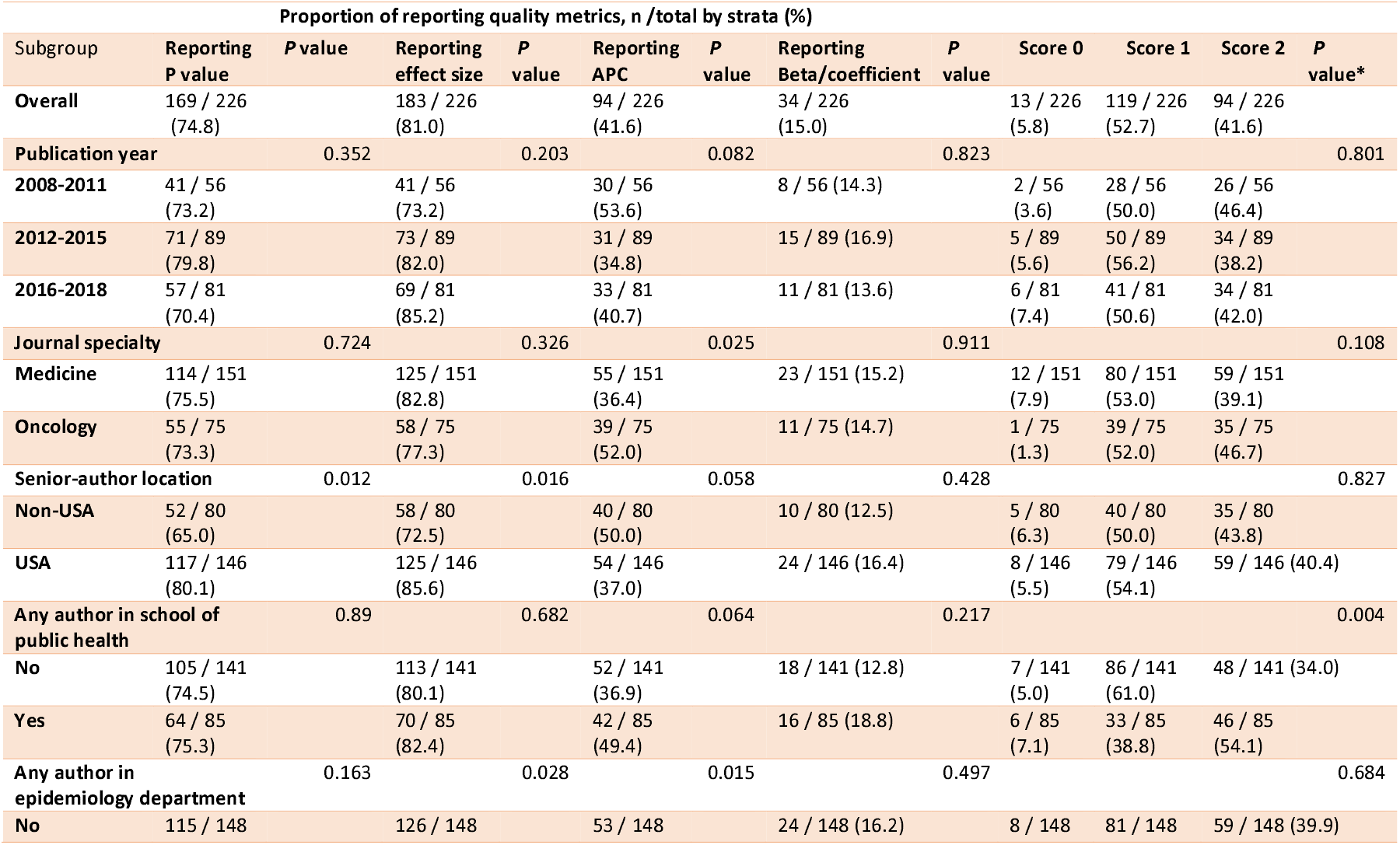
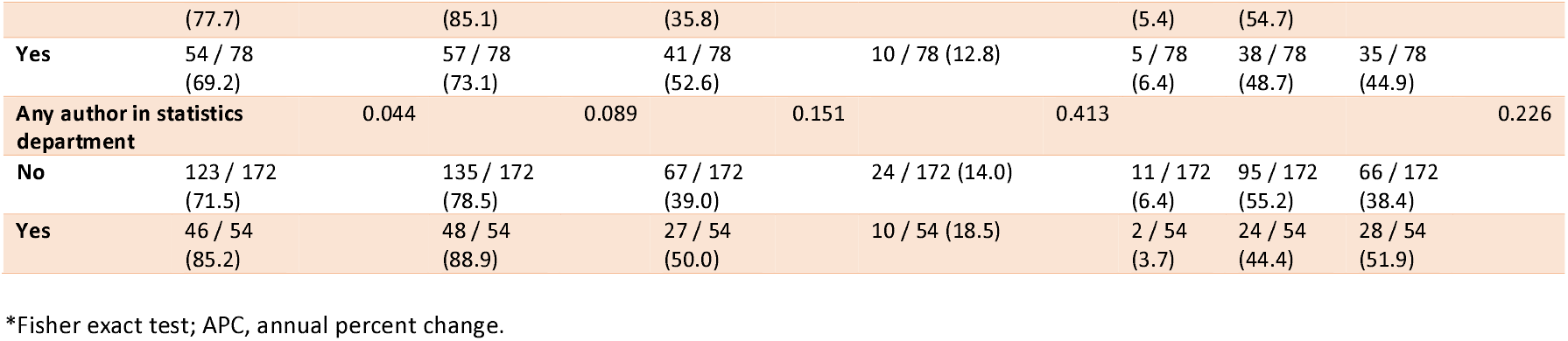
Reporting quality of the trend analyses with linear model published in leading medicine and oncology journals, 2008-2018.

**Table 2.**
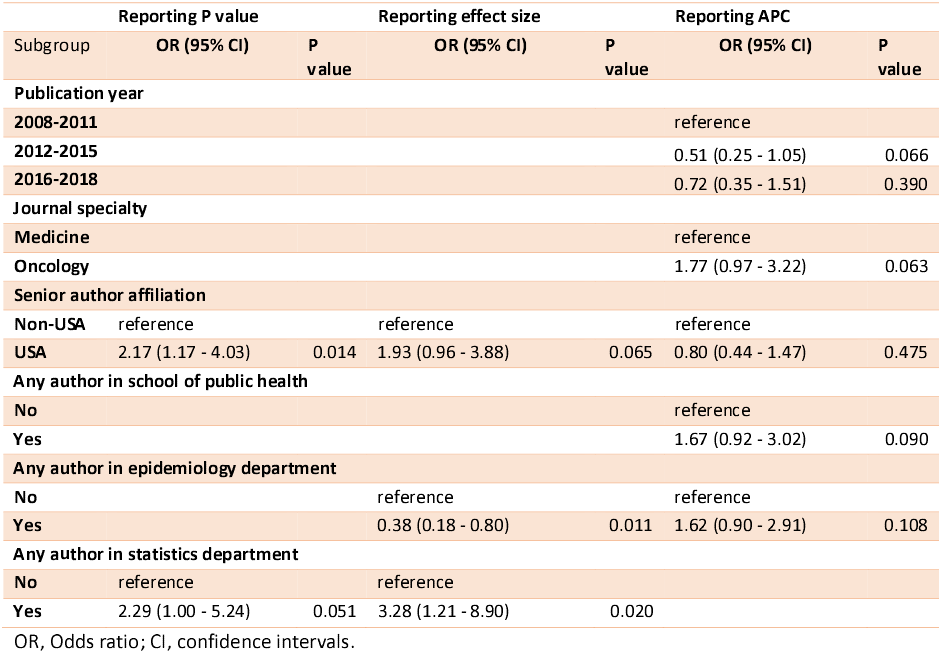
Multivariable models showing factors associated with reporting quality metrics of the trend analyses with linear model published in leading medicine and oncology journals, 2008-2018.

Reporting quality of the included trend analyses was overall good. This is consistent with reported increasing quality in clinical trials.^9^ However, several reporting-quality issues are concerning. Reporting p-value or effect-size did not change with publication years, despite the recommendation on using effect-size.^16 17^ The underlying reasons are worth additional investigation.

Moreover, Non-U.S. senior authors less frequently reported p-value or effect-size than U.S. ones, which warrants more research and training. Furthermore, more than half of the trend analyses using linear model did not report p-value/effect-size, slope/beta/APC or either, which is not consistent with recommendations.^15^ Without these metrics, quantification and comparison of linear models would be difficult or impossible. Therefore, the scientific rigor in these papers is significantly reduced.

In addition, we show that involvement of statistics department appears to link to more reporting effect sizes of oncological trend analyses, while that of epidemiology department linked to fewer reporting effect sizes. The rigorous training in statistics and its departments may contribute to this finding. Indeed, involvement of biostatisticians or epidemiologists was associated with higher methodological quality,^7^ higher acceptance rates,^20^ and shorter time to publication.^21^ Thus, it has also been advocated to include statisticians in more clinical research and trials.^10 22 23^

The paradoxical finding that epidemiology department involvement was associated with less effect-size reporting (OR=0.38, 95% CI: 0.18-0.80, p=0.011) warrants careful interpretation. One possible explanation is that the emphasis on effect size reporting is relatively recent (2016 and 2019), ^16 17^ yet epidemiologists may historically prefer other metrics. Given our sole focus on trend analyses in this study, this finding may not be applicable to, but should be examined in, other types of epidemiological and clinical studies. Strikingly, few studies on reporting quality of clinical trials or longitudinal studies separated biostatisticians and epidemiologists.^7 20 21^ Thus, future works are warranted to fill this knowledge gap.

There are few studies focused on reporting quality of trend analyses, while there are many on that of clinical trials or epidemiological studies.^7 9 11 12 24^ Some experiences and data in clinical trials and trend analyses may be applicable to each other. First, development and implementation of reporting guidelines appear to improve reporting quality of randomized clinical trials.^24^ However, there are no official reporting guidelines on trend analyses which may be needed. We thus recommend developing and implementing reporting guidelines on trend analyses, beyond recommendations.^15^ Second, librarians and information specialists did not impact reporting quality of systematic reviews but have helped with the journal review process.^25^ It will be interesting to understand whether librarians and information specialists will help improve reporting quality of trend analyses. Third, our findings seem to call for journal editors and peer reviewers to more rigorously enforce reporting standards and improve reporting quality. Indeed, one of the recommendations was published by a leading medical journal.^17^ Publishers and professional societies may also be more actively involved in enforcing reporting standards of trend analyses. Finally, our data seem to call for more involvement of statisticians in trend analyses in oncology research, probably also medical research at large. It is concerning that involvement of epidemiological department was associated with less reporting of effect size. More works are needed to confirm our findings and understand the reason. It is noteworthy that involvement of either department was not linked to better reporting scores.

This study has several limitations. First, our search strategy using only ‘trend/trends’ in titles may have missed relevant analyses using alternative terminology (e.g., ‘temporal changes,’ ‘secular patterns’). Second, we focused on high-impact journals, limiting generalizability to the broader literature. However, it is possible that journals of intermediate or low impact may publish trend analyses with worse quality than those of high impact. Future works are warranted to test this hypothesis. Third, we did not include any trend analyses published after 2018. These recent works are interesting due to the surges in publication numbers during and after the COVID-19 pandemic, ^18,19^ but may require a longer timeframe and more data than currently available data to reliably examine the biases and impact associated with the COVID-19 pandemic. Otherwise, the conclusions may be inaccurate, if not misleading. Third, our title-based search may have missed some relevant articles, that mentioned trend analyses only in the abstract or text. Although these may be a small portion of truly relevant articles, this limitation might affect the generalizability of the findings. Finally, the used reporting-quality metrics may not be applicable to all reports. Indeed, some trends are difficult to accurately model with a single algorithm, and thus may not report some of the quality metrics.

In summary, the reporting quality of trend analyses in leading medicine and oncology journals could be further improved. We thus call for more research and awareness of reporting-quality in trend analyses in oncology research and beyond.

## Author Contributions

Drs Zhang and Wang had full access to all of the data in the study and takes responsibility for the integrity of the data and the accuracy of the data analysis. They equally supervised the works, and share the senior authorship.

Concept and design

Yuan, Wang, Zhang.

Acquisition, analysis, or interpretation of data

All authors. Drafting of the manuscript: Yuan.

Critical revision of the manuscript for important intellectual content

All authors. Statistical analysis: Yuan, Deng, Zhang.

Supervision

Zhang, Wang.

