## Supplementary figures and images for "Reporting quality of trend analyses published in leading medicine and oncology journals during 2008-2018"

### Suppl Figure

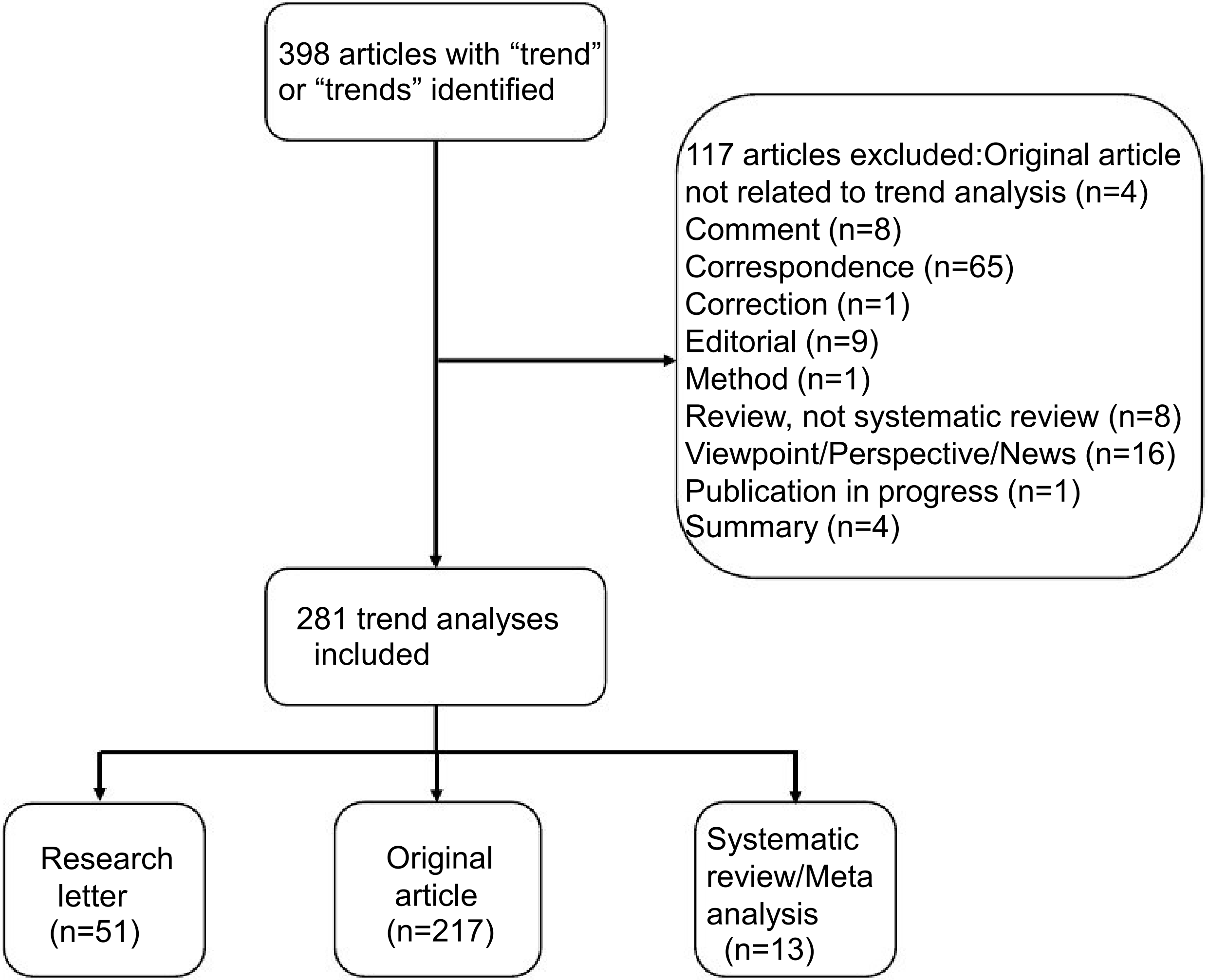
